# Statistical thermodynamics based design principles into the temperature induced fold switching of a metamorphic protein

**DOI:** 10.1101/2025.10.10.679401

**Authors:** Sridip Parui, Anand Srivastava

## Abstract

Fold-switching metamorphic protein sequences defy the classical “one sequence – one fold” paradigm. The ability of metamorphic proteins to reversibly switch between distinct folds make them attractive de novo protein engineering candidates since they can function as environment-sensitive molecular switches. However, the underlying thermodynamic design principles that drive their fold-switching behaviour is poorly understood. Gaining insights into the molecular driving forces leading to fold switching behavior is crucial for the rational design of new metamorphic proteins based molecular switches, sensors and stimulus responsive nanomaterials. In this study, we perform a detailed thermodynamic analysis of a designed fold-switching protein [*PNAS* 2023, *120* (4), e2215418120] ^1^ that transitions between a 3*α* fold and *α*/*β* fold upon changes in temperature. We use an efficient advanced sampling molecular simulation based free energy calculation approach called Confine-Desolvate-Convert-Solvate-Release (CDCSR), which subjects the protein through the complete range of thermodynamic cycle and deconvolutes the enthalpic and entropic driving forces at each stage of the cycle. We find that while 3*α* fold is stabilized at low temperatures by enthalpic contributions from favorable water-water and protein-water interactions, the transition to the *α*/*β* fold at high temperatures is driven by the gain of entropy from the release of ordered water molecules surrounding the 3*α* fold. Our study elucidates the molecular driving forces governing temperature-induced fold-switching behavior and provides a rigorous statistical thermodynamic framework that can help in the design and engineering of future synthetic and functional metamorphic proteins.

**Significance:** Recent success in the de novo protein design for molecular function is powered by the transformative AI methods and interpreted using the classical wisdom from the Physics-based modelling approaches. Stimuli-sensitive fold switching proteins that can take multiples shapes based on environmental factors are the next frontier in de novo protein engineering. In this work, we unravel the thermodynamic driving forces behind a recently designed temperature-sensitive artificial protein and provide a Physics-based framework to understand the design principles leading to fold switching behavior. Our work compliments the ongoing AI methods that are being explored to unravel the hidden evolutionary embeddings inherent in fold-switching proteins and also highlights the importance of thinking in terms of entropy-based design principles for natural system.

## Introduction

For decades, it was widely believed that a protein’s sequence leads to a single native fold. ^2^ However, recent discoveries ^1,3–6^ have revealed the existence of metamorphic proteins, which can adopt multiple distinct conformations without substantial changes in their amino acid sequences. This ability to switch between different conformations allows metamorphic proteins to exhibit diverse functionalities.^3,4,7–9^ The conformational transitions, however, are not spontaneous; they are triggered by environmental factors such as temperature, pH, salt concentration, or ligand binding.

Although only a handful of metamorphic proteins have been identified to date, there is growing consensus that they may be more prevalent than previously thought. The “Janus” nature of structural and functional duality of metamorphic proteins has therefore spurred significant interest in the prediction of their sequences ^12,13^ and design.^1,14–19^ However, current design strategies do not have a well-defined concept based prescription and largely depend on trail and error as well as the "intuition" of domain experts. Developing rule-based approaches requires a deeper understanding of how environmental factors influence protein conformations and the specific thermodynamic forces that drive fold switching. Understanding the driving forces behind fold switching requires determining the free energy difference (ΔG) between two conformations, as free energy quantifies stability. Since population serves as a proxy for free energy, previous NMR studies^1,10,20^ have examined the populations of two conformations under varying temperatures and salt concentrations. These studies reveal which structure is more stable under specific conditions. However, identifying the exact factors contributing to differential stability is challenging due to the limited molecular precision of experiments and associated technical constraints. These limitations often prevent the direct assessment of the distinct thermodynamic roles of the protein and solvent.

Physics-based molecular dynamics (MD) simulations can complement experimental data by providing a detailed, atomistic description of protein structures in ensemble form.^21–23^ However, conformational transformations often occur on timescales that are sometimes practically unattainable in traditional MD and thus making it computationally costly. Efficient exploration of conformational dynamics often relies on enhanced sampling MD methods, which employes biasing potentials along predefined reaction coordinates. Yet, identifying appropriate reaction coordinates that are both easy to sample and relevant to fold switching remains a significant challenge, particularly for larger proteins.

In this study, we employ a reaction coordinate-free MD approach, utilizing a thermodynamic cycle we call Confine-Desolvate-Convert-Solvate-Release (CDCSR), to compute the free energy differences between two protein folds and investigate the driving forces behind fold switching. We applied this method to a recently designed metamorphic protein, which undergoes a temperature-induced fold switch, as reported by Orban and colleagues. ^1^ This protein exhibits two conformations: the 3-*α* fold, which predominates at low temperatures, and the *α*/*β* fold, which is more stable at higher temperatures. To understand the thermodynamic underpinnings of this fold-switching behavior, we provide a detailed analysis that decouples the contributions of various components in the system. Using the CDCSR method, we show that the 3-*α* fold is stabilized at low temperatures through favorable water-water and protein-water interactions, while the *α*/*β* fold is favored at higher temperatures due to an increase in water entropy. These insights contribute to a deeper understanding of the design principles of metamorphic proteins and may guide future efforts in developing proteins with switchable conformational states.

## Methods and Material

### Confine–Desolvate–Convert–Solvate–Release (CDCSR): A thermodynamic cycle for free energy change from A to B

To estimate the free energy of conformational change, reaction coordinate-based molecular dynamics (MD) simulations are commonly employed.^24–26^ These convert conformation A to conformation B through a series of physically meaningful intermediate states. However, traversing these intermediates can be computationally challenging. But free energy being a state function depends only on the initial state A and final state B, regardless of the pathway taken. Previous studies show that a thermodynamic cycle that involves unphysical routes for the conversion following confinement simulation approaches and principles of statiscal mechanics can provide efficient estimation of free energy of fold switching (Δ*F*_*AB*_).^27-30^ Roy et al^28^ implemented such thermo cycle named Confine-Convert-Release (CCR) using implicit model on AMBER software suite. To explicitly account for solvent effects, we extend this framework and propose a five-step thermodynamic cycle (see Figure 1 1), termed Confine–Desolvate–Convert–Solvate–Release (CDCSR). This cycle is implemented using GRO-MACS^31–34^ with an explicit solvent model. The detailed procedure for each step of the CDCSR method is given below.

**Figure 1:**
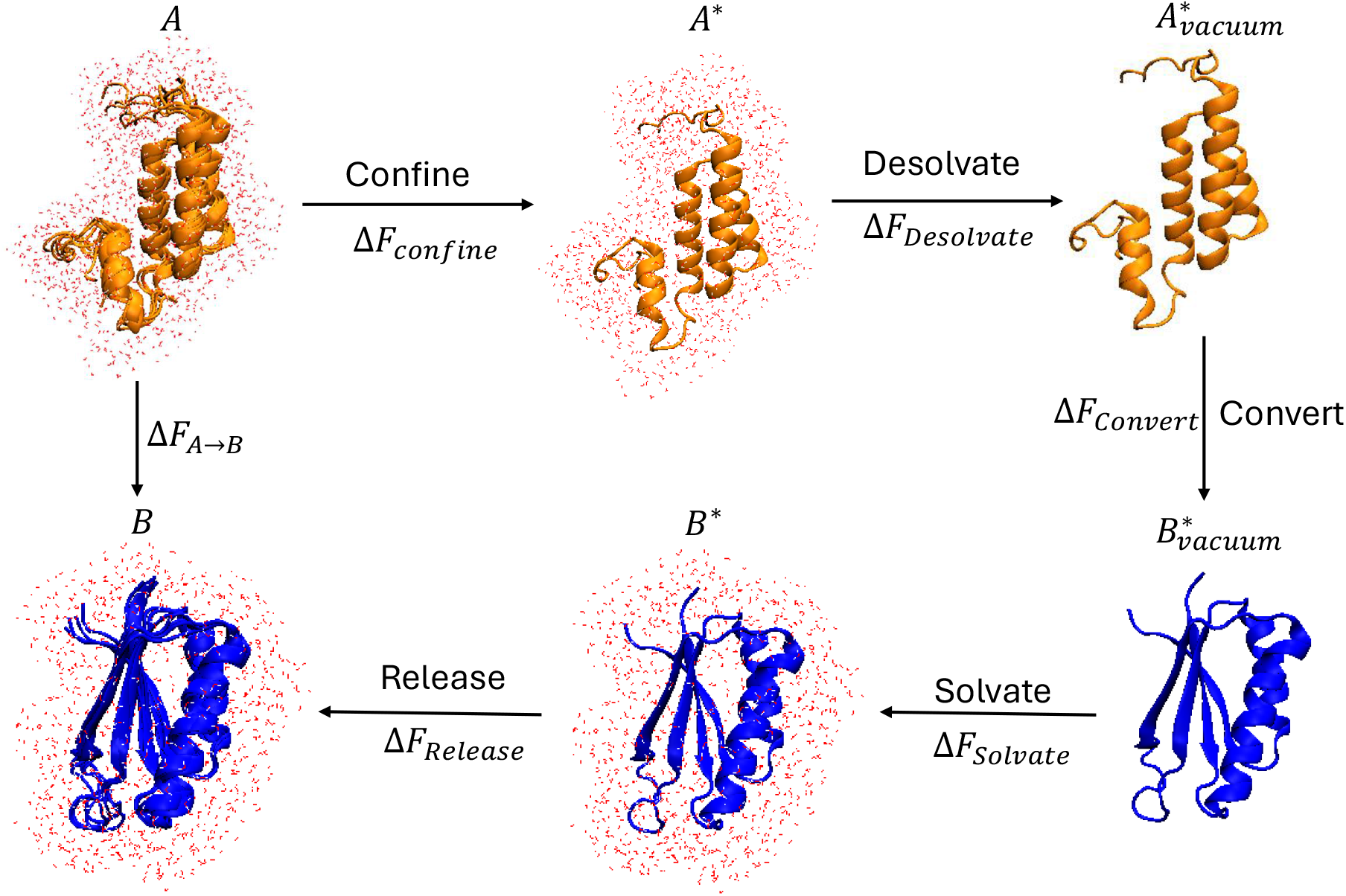
Free energy of fold switching from *α* to *α/β* using CDCSR. The thermodynamic cycle consists of five steps. (i) The conformational dynamics of state A^∗^ are confined to a single microstate-like reference state, *A*^∗^, by applying positional restraints. (ii) *A*^∗^ is desolvated to obtain 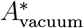. (iii) 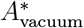 is converted into 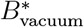, a corresponding single microstate-like reference for B in vacuum. This is a not a physical transformation; the free energy change is obtained as the difference between the individually computed free energies of 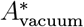 and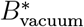, each evaluated from their potential energies and vibrational entropies. (iv) 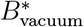 is solvated to obtain *B*^∗^. (v) Finally, the position restraints on *B*^∗^ are released to get the original conformational ensemble of B.

#### Confine

The first step involves confining the initial state A which represents ensemble of conformations within its native basin due to the thermal motion. This conformational flexibility is gradually frozen into a reference microstate, *A*^∗^ by imposing position restraints on all atoms of the protein. The strength of these restraints is increased progressively until *A*^∗^ represents a nearly single microstate. The free energy for this step is confinement free energy (Δ*F*_confine_) which also is a measure of conformational entropy of state A. The theoretical framework for this step is detailed in the Appendix A.

#### Desolvate

Next, the reference state *A*^∗^ is desolvated to obtain its vacuum equivalent, 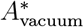. This can be done by gradually turning off the electrostatic and van der Waals potentials of water, while maintaining the restraints on the protein atoms. The free energy change for this process (Δ*F*_desolvate_) corresponds to the solvation free energy of *A*^∗^ with the opposite sign, i.e., Δ*F*_desolvate_ = −Δ*F*_solvate_. Further details are provided in the Appendix B.

#### Convert

This step involves the conversion of 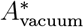 to *B*^∗^vacuum, both of which are highly restrained, near-static reference states in vacuum. However this is not an actual conformational transformation in simulation. Here, we compute the free energies of these two states individually. Their difference gives the conversion free energy, Δ*F* convert. Since these are tightly restrained systems, their free energy includes only potential energy and vibrational entropy which is estimated using the quasi-harmonic approximation (QHA). The use of QHA is justified due to use of strong restraints on all the atoms of the protein. A detailed explanation is provided in the Appendix C.

#### Solvate

The vacuum state, 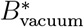 is then solvated to obtain *B*^∗^. This is done by gradually turning on electrostatic and van der Waals potentials of water. The resulting free energy change, Δ*F*_solvate_, corresponds to the solvation free energy of *B*^∗^.

#### Release

Finally, the restraints on *B*^∗^ are gradually removed to restore its native ensemble character resulting in the final state B. The free energy change for the release (Δ*F*_release_) is basically the confinement free energy of B (Δ*F*_confine_) with opposite sign, i.e., Δ*F*_confine_ = −Δ*F*_release_. The confine and release steps are same but opposite in direction, as are the desolvate and solvate steps.

The overall free energy of the conformational transition from A to B is therefore,

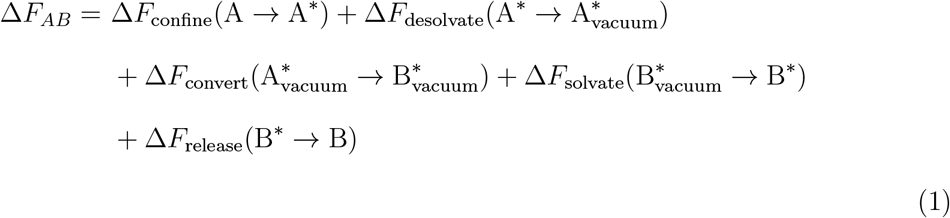

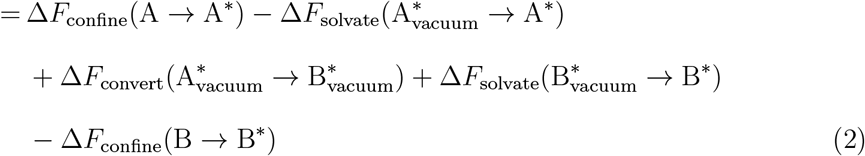

### Simulation details

#### System Preparation

We selected the metamorphic sequence Sa1_V90T from Orban and coworkers.^1^ The initial structures include a 3*α* conformation obtained from PDB-IHM (PDB ID: 9A20; DOI: 10.2210/pdb9a20/pdb) and *α/β* conformation from RCSB PDB (PDB ID: 8E6Y; DOI: 10.2210/pdb8E6Y/pdb). Simulations were performed on GROMACS 2024.2^31–34^ patched with PLUMED 2.9.2^35–37^ packages. We have selected the CHARMM36^38,39^ force field with the TIP3P*^40^ water model. For comparison, we also tested the Amber ff99SB-ILDN force field^41^ with TIP3P^40^ water and Amber-disp,^42^ but CHARMM36 was found to be best among three for our study (see SI for details). Both structures were solvated in a cubic box (7.5 nm x 7.5 nm x 7.5 nm) that contains 13,812 water molecules and two chloride ions to neutralize the system. Energy minimization of the solvated systems was performed using the steepest descent method.

#### Equilibrium Simulation

The minimized structure was equilibrated in two stages conducted at five different temperatures: 260 K, 280 K, 300 K, 320 K, and 340 K. First, an NVT equilibration was done by restraining the protein’s heavy atoms with a force constant of ∼240 kcal-mol^−1^-nm^−2^ for 1 ns. The V-rescale thermostat, a modified Berendsen thermostat, was used to maintain the temperature. The final configuration from the NVT equilibration was then used as the starting point for a position-restrained NPT equilibration performed for 10 ns. Pressure was maintained at 1 bar using the Parrinello-Rahman barostat. Following the NPT equilibration, a final steepest descent energy minimization was performed on the resulting configurations. These minimized structures served as the input starting structures for CDCSR simulations at their respective temperatures.

#### Confine and Release

In the confinement steps, both *α* and *α/β* conformers are gradually restricted to a single microstate-like situation. This was done by applying position restraints on all protein atoms. In the free state, the conformer has negligible restraint, with a force constant of 3.6 × 10^−6^ kcal·mol^−1^·nm^−2^. The force constant was progressively increased to approximately 3830 kcal·mol^−1^·nm^−2^ across 16 replicas. To enhance conformational sampling in the free state, Hamiltonian Replica Exchange Molecular Dynamics (HREMD),^43,44^ as implemented in PLUMED patched with GROMACS, was employed. For HREMD, an additional 16 replicas were included, where potentials were scaled with the *λ*) ranging from 1.0 to 0.59. Further details are provided in Table S1. Confinement simulations were performed under the NVT ensemble where temperature was maintained by the V-rescale thermostat.^45^ The leap-frog algorithm was used to integrate the Newton’s equations of motion with a time step of 2 fs. Long-range electrostatic interactions were calculated using the Particle Mesh Ewald (PME) method with a cutoff of 1.0 nm and a grid spacing of 0.16 nm. Van der Waals interactions were also calculated with a cutoff of 1.0 nm. Each replica was simulated for 10 ns. First 100 ps was discarded for analysis. For calculation of confinement free energy integration recipe given by Karplus and coworkers was used. ^27^

#### Desolvate and Solvate

In these steps, both *α* and *α/β* still remain fully confined with a force constant of approximately 3830 kcal·mol^−1^·nm^−2^. The free energy feature in GROMACS was utilized to gradually reduce the interaction potentials of water to zero to mimick the desolvation of the protein into a vacuum. First, the Coulombic potential of water was scaled down linearly using a scaling factor *λ*. Afterward, the van der Waals potential of water was scaled in a similar manner. This two-step scaling process was implemented to prevent the collapse of interacting atoms. The scaling factors ranged from 0 to 1 and were distributed over 16 replicas. See Table S2 for details. A soft-core potential was applied for the van der Waals interactions to avoid singularities, with *α* = 0.5 and *σ* = 0.3 nm. The interaction potential derivative with respect to the scaling factor, 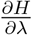, was printed every 100 steps, equivalent to 200 fs, as the integration time step was set to 2 fs using the leap-frog stochastic dynamics (SD) integrator. Temperature coupling was implicitly handled by the SD integrator. The total simulation time per replica was 4 ns, with the first 100 ps discarded for analysis. Free energy for desolvation (or solvation) was calculated by using `gmx bar` command in GROMACS.

#### Convert

In this step, ensemble average of the potential energy for *α/β* was first estimated. In the vacuum state, the protein also has vibrational entropy. Rotational and translational modes are excluded due to the restraints applied to the protein. The free energy of *α/β* in vacuum was determined as the sum of the average potential energy and the vibrational entropy contribution (−*TS*_vib_). To calculate the vibrational entropy, the eigenvalues for each of the 4635 modes (the protein contains a total of 1545 atoms) were determined using the `gmx covar` command. The vibrational entropy was then calculated by skipping the rotational and translational modes using the `gmx anaeig` command with consideration of quasi harmonic approximation (QSA). Similarly, the free energy for the *α* state in vacuum was computed. The difference between the free energies of the *α/β* and *α* states provided the conversion free energy.

## Results and Discussion

### CDCSR correctly predicts 3*α* stability at low temperatures and *α/β* stability at high temperatures

To investigate the driving forces behind fold switching, we calculated the free energy change (Δ*F*) for the transition from the 3*α* to the *α/β* conformation at different temperatures (260 K, 280 K, 300 K, 320 K, and 340 K). The plot of Δ*F*_3*α*→*α/β*_ versus temperature (Figure 2A) shows that Δ*F* is positive at low temperatures, indicating that the 3*α* fold is more stable at low temperatures. As temperature increases, Δ*F* decreases and eventually becomes negative, suggesting that the *α/β* fold is favored at higher temperatures. Although the simulation results deviate quantitatively (,^1^ the overall trend is consistent with experimental observations. It is to be noted that the error bars on Δ*F* are larger at intermediate temperatures (300 K and 320 K). In this range, the magnitude of Δ*F* is reduced and becomes zero at transition temperature, suggesting that both 3*α* and *α/β* folds coexist in equilibrium. Presence of both folds within this temperature range leads to higher uncertainty in free energy estimation.

**Figure 2:**
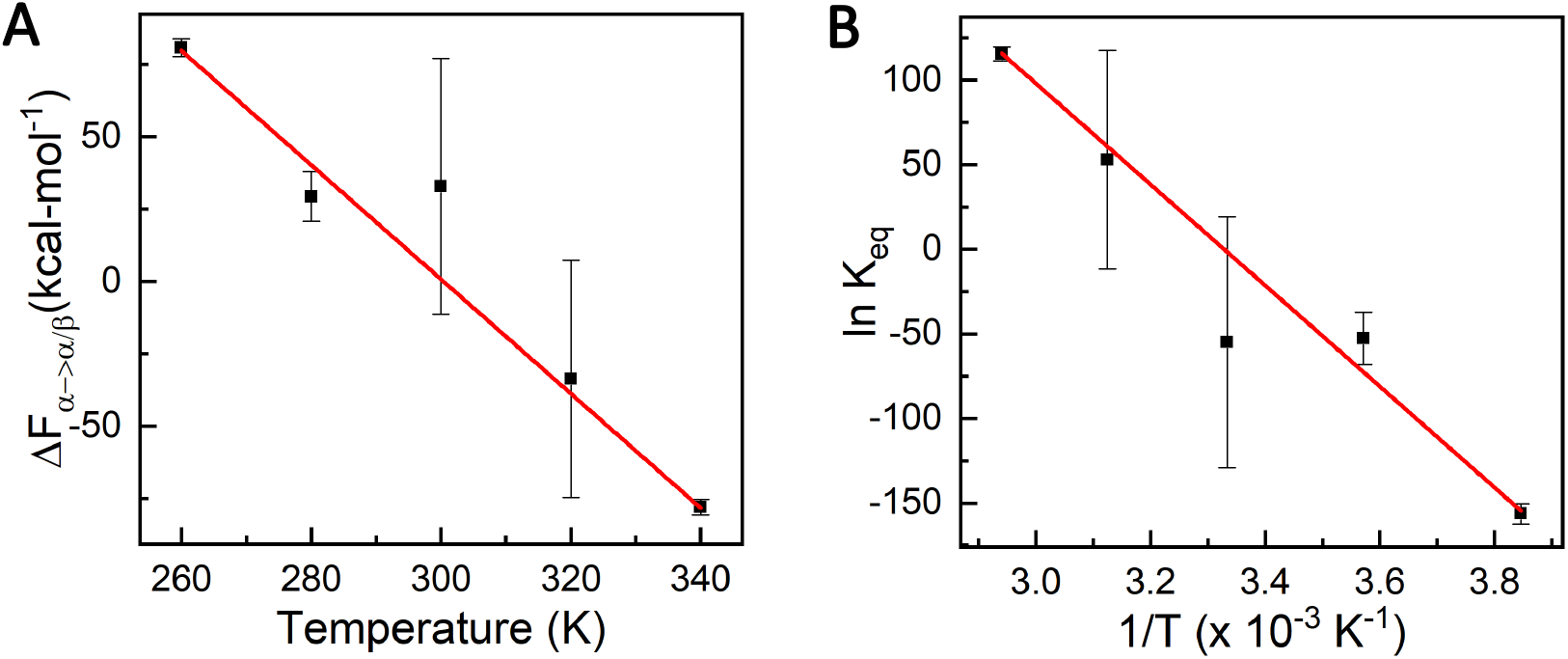
Temperature-dependent relative stability of 3*α* and *α/β* conformers. (A) Free energy difference for the conversion (Δ*F*_*α*→*α/β*_) as a function of temperature. At low temperature, Δ*F* is positive, indicating that 3*α* is more stable. With increasing temperature, Δ*F* decreases and becomes negative, showing that *α/β* becomes favored at high temperature. (B) van’t Hoff plot (ln *K*_*eq*_ vs. 1*/T*), where *K*_*eq*_ is the equilibrium constant for the conversion and T is temperature. The negative slope indicates endothermic nature of the process (Δ*H*° *>* 0), while the positive intercept indicates an entropy gain (Δ*S*° *>* 0) for the 3*α* → *α/β* transition.

### van’t Hoff analysis reveals endothermic nature of 3*α* to *α/β* transition driven by entropy gain

To determine whether the transition is endothermic or exothermic, we obtained van’t Hoff plot (ln *K*_*eq*_ vs. 1*/T*) (Figure 2B). The van’t Hoff relation is given by:

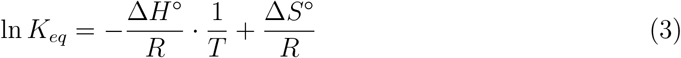

where *K*_*eq*_ is the equilibrium constant which is related to the free energy change as:

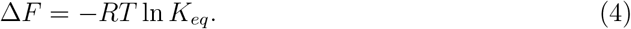

Here, *T* is temperature, Δ*H*° is the standard enthalpy change, and Δ*S*° is the standard entropy change. The van’t Hoff plot fits well to a straight line with *R*^2^ = 0.99. The negative slope (−Δ*H*°*/R*) indicates an endothermic nature (Δ*H*° *>* 0) of the 3*α* to the *α/β* conversion. The positive intercept (Δ*S*°*/R*) corresponds to positive entropy change. While the estimated values (Δ*H*° ≈ 590 kcal mol^−1^, Δ*S*° ≈ 1.9 kcal mol^−1^K^−1^) are much larger than experimental data, they still provide meaningful qualitative insight. The discrepancy possibly arises from the limitations in force field accuracy for metamorphic proteins. However, the observed endothermicity and positive entropy change are in agreement with experimental findings.

### 3*α* to *α/β* transformation is driven by increase in water entropy

Van’t Hoff analysis provides the standard enthalpy (Δ*H*°) and entropy (Δ*S*°) changes. However, to understand the precise thermodynamic driving forces, it is necessary to evaluate actual Δ*H* and Δ*S* at different temperatures. The entropy change (−Δ*S*) at a given temperature *T* can be estimated using the finite difference method with central difference approximation:

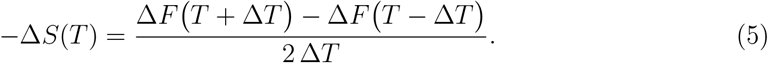

In this study, entropy was calculated at 280 K, 300 K, and 320 K with Δ*T* = 20 K. Due to the central difference scheme, entropy values at the end points (260 K and 340 K) could not be obtained. The enthalpy change (Δ*H*) at temperature *T* was then determined as:

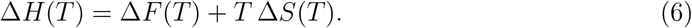

Figure 3A shows the decomposition of the free energy change (Δ*F*) into enthalpic (Δ*H*) and entropic (−*T* Δ*S*) contributions. The results clearly demonstrate that −*T* Δ*S* for the 3*α* → *α/β* conversion is negative and becomes increasingly negative with temperature. This indicates that the transformation to *α/β* is entropically driven, whereas the 3*α* state is stabilized by enthalpy.

**Figure 3:**
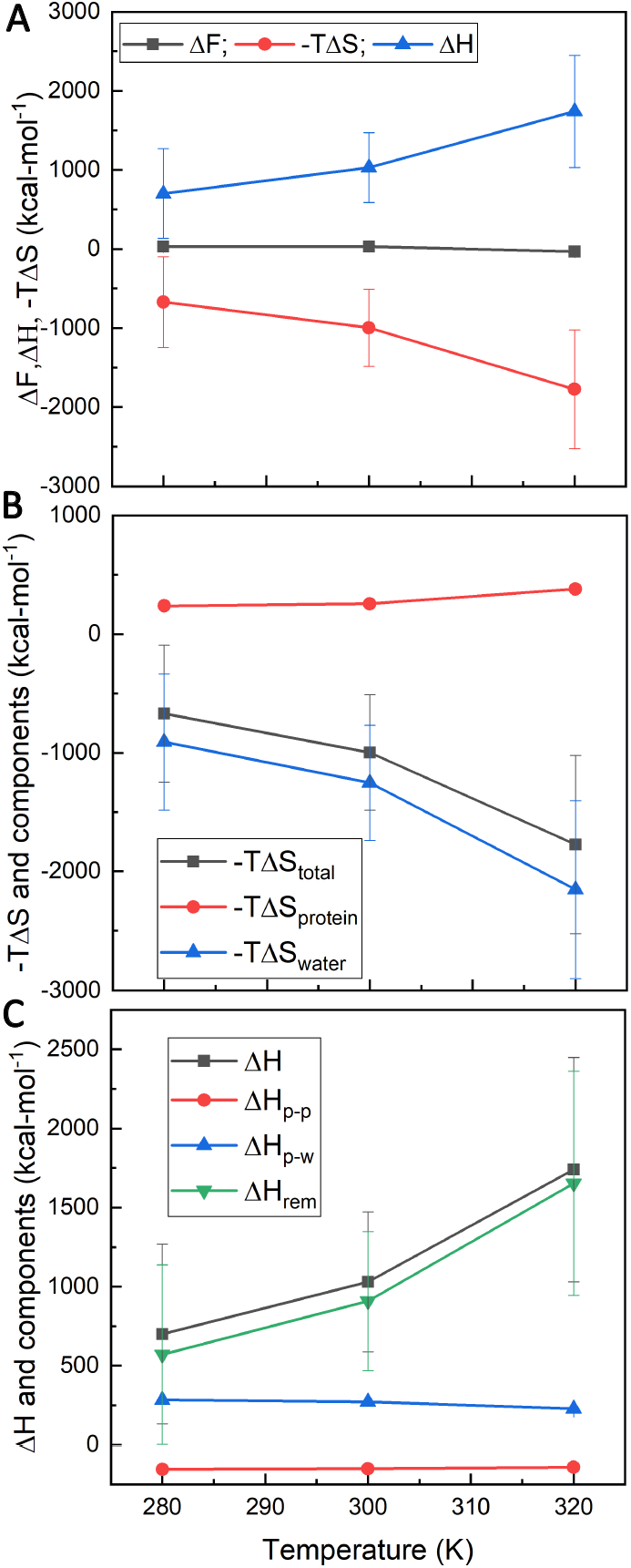
Decomposition of free energy change (Δ*F*) into enthalpic and entropic contributions for the 3*α* → *α/β* transition at different temperatures. (A) Decomposition of Δ*F* into entropy (−*T* Δ*S*) and enthalpy (Δ*H*) shows that the favorable entropic contribution for this transformation increases with temperature, while the associated enthalpic contribution is unfavorable and becomes more unfavourable at higher temperatures. This means that the reverse process, i.e., formation of 3*α*, is enthalpically favored. (B) Decomposition of the entropy contribution into protein and water shows that the favorable entropy driving the formation of *α/β* is mainly from water. (C) Further decomposition of Δ*H* into protein-protein (Δ*H*_*p*−*p*_), protein-water (Δ*H*_*p*−*w*_), and residual enthalpy (Δ*H*_*rem*_, dominated by exclusively water-water interactions) shows that the stability of 3*α* is primarily due to favorable water-water interactions and protein-water interactions to lesser extent.

Entropy has two main components: protein entropy (Δ*S*_protein_) and water entropy (Δ*S*_water_):

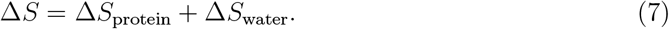

The protein entropy contribution includes confinement free energy (-Δ*F*_confine_), which corresponds to conformational entropy, and vibrational entropy (Δ*S*_vib_):

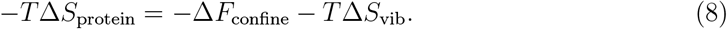

Accordingly, the water entropy contribution is given by:

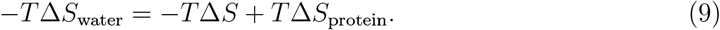

Figure 3B shows the decomposition of entropy into protein and water contributions. The results indicate that while the 3*α* → *α/β* transition is associated with a positive protein entropy contribution (−*T* Δ*S*_protein_), the water entropy contribution (−*T* Δ*S*_water_) is negative and becomes more negative at higher temperatures. Thus, the overall entropic stabilization of the *α/β* fold originates predominantly from water entropy.

### Water-water and protein-water interactions favor 3*α* **over** *α/β*

To gain deeper insight, the total enthalpy change can be decomposed into protein–protein (Δ*H*_p-p_), protein-water (Δ*H*_p-w_), and water-water contributions (Δ*H*_*w*−*w*_). Δ*H*_p-p_ and Δ*H*_p-w_ can be directly obtained from the simulation. The remaining enthalpy change, (Δ*H*_rem_) is,

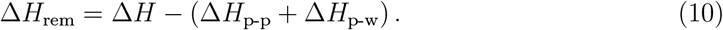

Δ*H*_rem_ is dominated by water-water interactions, as other contributions such as mechanical pressure volume work are negligible. Therefore,

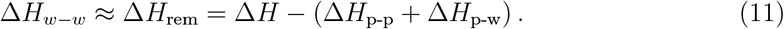

It is to be noted that direct estimation of Δ*H*_*w*−*w*_ from the simulation gives rise to high uncertainty as the fluctuation in water-water interactions is quite high.

Figure 3C shows the decomposition of Δ*H* for the 3*α* → *α/β* transformation. In this case, both Δ*H*_rem_ and Δ*H*_p-w_ are positive, whereas Δ*H*_p-p_ is negative. This indicates that the reverse transformation (*α/β* → 3*α*) is largely stabilized by water-water interactions, with protein-water interactions contributing to a lesser degree.

### Residue-wise analysis reveals stronger water–water interactions at solvent-exposed terminals in 3*α*

To identify which protein segments or residues contribute most to the enthalpic stabilization of the 3*α* state, we computed the residue-wise water–water interaction enthalpy change (Δ*H*_*w*−*w*_) for 3*α* → *α/β* transformation (Figure 4A). Only water molecules within the first solvation shell (Figure S5) of each residue were considered, and the calculation included only nonbonded interactions among water molecules.

**Figure 4:**
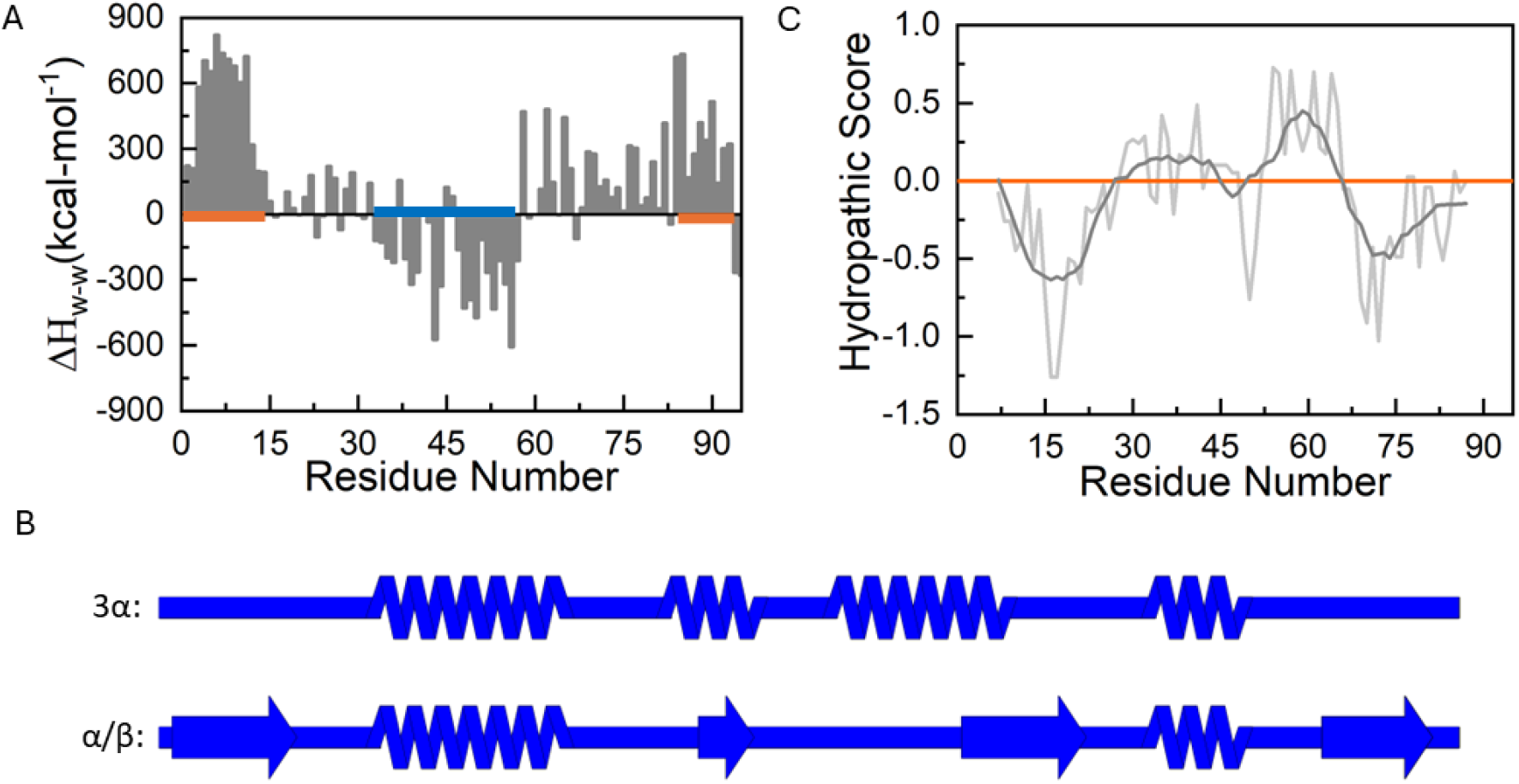
Residue-level analysis of factors governing the stability of 3*α* and *α/β*. (A) Change in water–water interactions within the first solvation shell of each residue for the 3*α* to *α/β* transformation. The terminal regions (residues 1–15 and 83–94) show a continuous stretch of positive Δ*H*, indicating stronger water-water interactions in 3*α* compared to *α/β*. (B) Secondary structure comparison between 3*α* and *α/β* highlights that the terminal segments of 3*α* are unstructured. The diagram was generated using SSDraw software.^46^ (C) Kyte–Doolittle hydropathy profile^47^ along the sequence, obtained using the ExPASy server,^48^ which shows lower hydropathy scores at the terminal regions providing a possible rationale for the unstructured nature of the terminals in 3*α*.

Figure 4A shows that the terminal regions, particularly the N-terminal segment (residues 1–15, left orange trace) and, to a lesser extent, the C-terminal segment (residues 83–94, right orange trace), have continuous stretch of positive Δ*H*_*w*−*w*_. This indicates that water-water interactions are stronger around these regions in the 3*α* form compared to the *α/β* form. Residue segment 60–80 also show positive Δ*H*_*w*−*w*_, but the effect is weaker and more irregular. Interestingly, while the terminal segments are associated with enhanced water-water interactions in 3*α*, the intermediate segment (residues 32–58, blue trace) shows stronger interactions in the *α/β* state.

Structural inspection reveals that the terminal regions are disordered and solvent-exposed in the 3*α*, but folded in the *α/β* form (Figure 2A, Figure 4B and Figure S6). Solvent exposure of the terminals in 3*α* explains the more favorable protein–water interactions relative to *α/β*. We also analyzed the hydropathy profile along the sequnce (Figure 4C). The results clearly show that the terminal segments have negative hydropathic score compared to the intermediate segment. Reduced hydrophobicity accounts for the disorderness and solvent exposure of the terminals in 3*α*. Since hydrophobic groups are believed to promote structural ordering of water, increased solvent exposure of terminal hydrophobic residues in 3*α* likely contributes to stronger water–water interactions and its enthalpic stabilization at low temperatures.

### Water structure and ordering drive stability of 3*α* **and** *α/β*

It is well established that the structure and dynamics of water play a crucial role in governing protein stability and conformational fluctuations. ^49–52^ To connect the thermodynamic analysis with water structure and dynamics, we investigated the hydrogen bond angle (OOH) distribution and oxygen-oxygen auto correlation function (Figure 5). Calculation of angle was done by considering oxygen–oxygen pairs within 0.4 nm, which is around the first minimum of the oxygen–oxygen radial distribution function of water (Figure S7). Previous study (^53^ have also used this 0.4 nm cutoff, as it provides an optimal balance between capturing solute-induced perturbations and unperturbed water structure.

**Figure 5:**
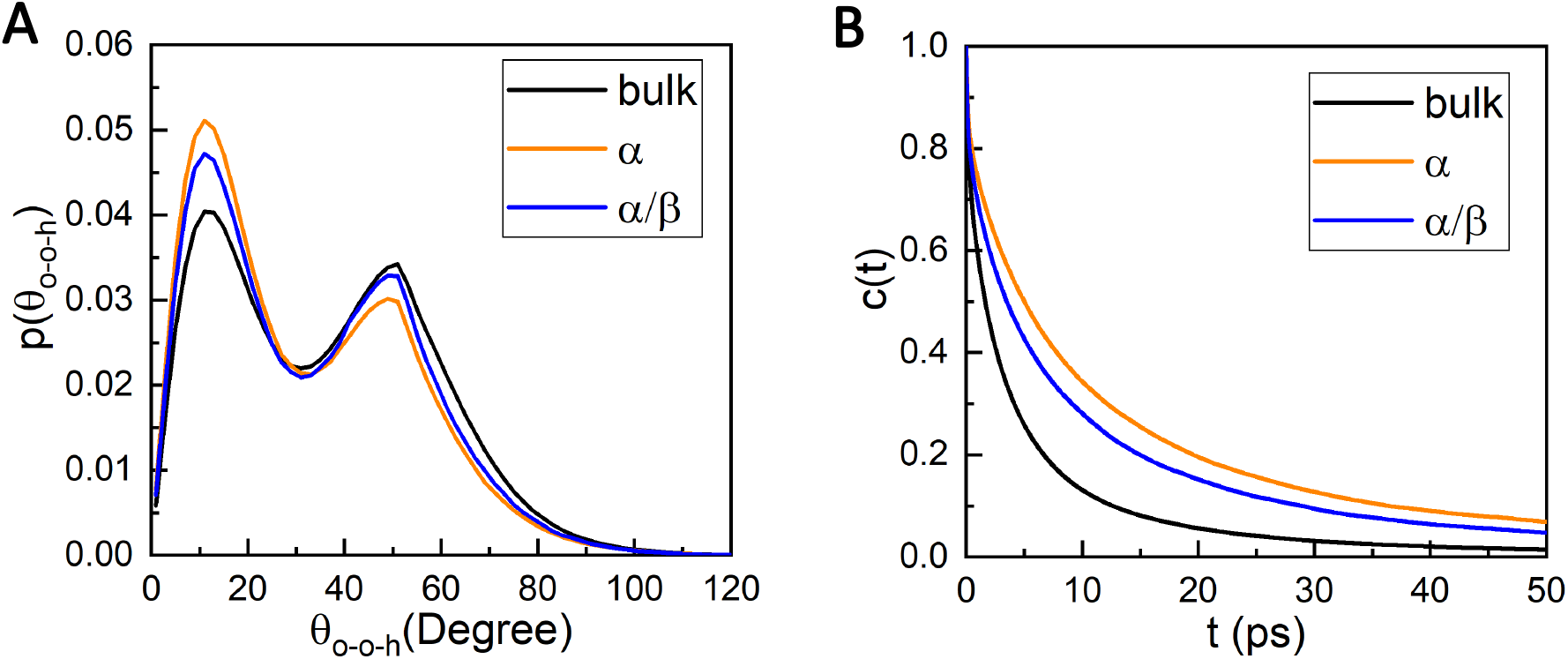
Hydrogen bond structure and dynamics of water. (A) O–O–H angle distributions for bulk water and water near the terminals of 3*α* and *α/β*. The distribution shows two peaks: a strong, linear hydrogen bond near ∼15#x00B0; and a weaker, bent bond near ∼50°. Water around the 3*α* terminal forms stronger hydrogen bonds compared to *α/β* and bulk. (B) Distance-based water–water autocorrelation functions in bulk and near *α* and *α/β*. Water near the 3*α* shows the slowest decay, which indicates the highest life time of water-water bonds among the three, particularly in the fast decay component.

Typical OOH distribution exhibits two peaks: one near 15° and another near 50°. The 15° peak corresponds to linear hydrogen bonds which indicates stronger hydrogen bonds. The 50° peak represents bent hydrogen bond which is associated with weaker interactions. We first analyzed water molecules surrounding the entire protein (Figure S8). Relative to bulk, water around both the 3*α* and *α/β* show higher population of stronger hydrogen bonds. Between the two, the *α* form has slightly higher population of linear hydrogen bonds. This trend is even more pronounced when water molecules near the terminal residues (Figure 5A) are considered. The enhanced ordering of water at the terminal explains why the 3*α* form is stabilized predominantly through water–water interactions.

To investigate water dynamics, we calculated the hydrogen bond autocorrelation function as,

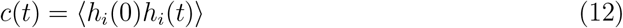

where *h*_*i*_(*t*) = {0, 1} for hydrogen bond *i* at time *t*; *h* = 1 if the hydrogen bond criterion is satisfied and *h* = 0 otherwise. Initially, we defined a hydrogen bond by an oxygen–oxygen distance below 0.4 nm and an OOH angle below 30°. Compared to bulk (Figure S9), the decay of the correlation function of hydrogen bonds in water surrounding both 3*α* and *α/β* is slower, which suggests longer lifetimes of hydrogen bonds near the proteins. This is consistent with the presence of stronger hydrogen bonds, as inferred from the OOH angle distribution. Interestingly, the hydrogen bond lifetimes are comparable for both conformers under this definition of hydrogen bond.

However, when we relaxed the definition and considered only the distance criterion in autocorrelation calculations, the decay rates of distance-based correlation functions differed appreciably (Figure 5B). In this case, water around the 3*α* conformer shows longer life-times than water near the *α/β* conformer. The higher lifetime suggests stronger and more persistent water–water interactions around 3*α*.

In a nutshell, the terminal regions of 3*α* are solvent-exposed, that results in hydrophobic residues accessible to water. Around these exposed hydrophobic groups, water forms strongly hydrogen-bonded, ordered networks compared to both *α/β* and bulk water. This ordering rationalizes the stability of the 3*α* state at low temperature through water–water interactions. But the 3*α* → *α/β* transition involves structural compaction and reduced solvent exposure of hydrophobic residues (Figure S10), which leads to decreased water ordering. This reduction contributes to positive entropy change from water, which favors the *α/β* state at higher temperatures. Thus, the stability of *α/β* at higher temperatures is driven by water entropy, while the stability of 3*α* at low temperature arises from favorable water structuring.

## Conclusions

In this study, we have demonstrated a computational framework for computing free energies of fold switching proteins and provided a detailed thermodynamic analysis of the temperature-induced fold switching in a designed metamorphic protein. Our results show that the stability of the 3*α* fold at low temperatures is enthalpically driven by favorable water-water and protein-water interactions, while the transition to the *α/β* fold at high temperatures is driven by an increase in water entropy. The thermodynamic insights gained from this work are crucial for understanding the molecular principles underlying fold switching and can help in the design of future metamorphic proteins. Further studies are needed to refine our computational approaches and explore additional factors such as pH, salt that may influence the fold switching behavior.

## Appendix A: Theory for confinement free energy

To compute the confinement free energy, we define two states: a *free state A* and a *reference confined state A*^∗^. In the reference state, all atoms of the protein are strongly position-restrained such that the potential energy function becomes

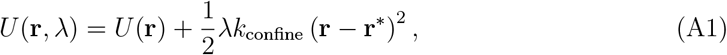

where *U* (**r**) is the original force field potential at atomic configuration **r**, *k*_confine_ is the harmonic force constant for the restraint employed in the reference state *A*^∗^, and **r**^∗^ is the atomic position vector in *A*^∗^. The scaling factor *λ* ∈ [0, 1] interpolates between the unrestrained free state (*λ* = 0) and the fully confined reference state (*λ* = 1).

Using thermodynamic integration, the confinement free energy for *A* to *A*^∗^ is expressed as

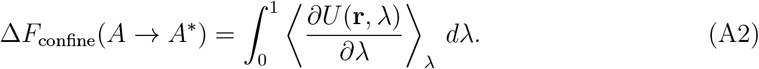

The derivative of the potential energy with respect to *λ* for a given configuration **r** is

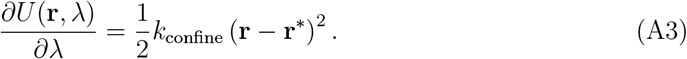

Substituting this into the integral gives

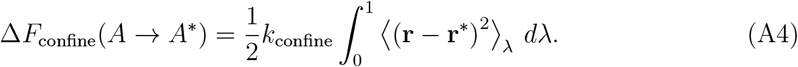

If *k* = *λk*_confine_, we can rewrite the integral in terms of force constant *k*:

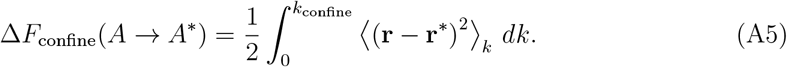

The root mean square deviation (RMSD) between a configuration **r** and the reference **r**^∗^ is defined as,

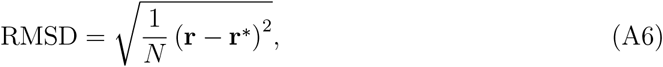

where *N* is the number of atoms in the protein. Therefore,

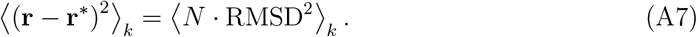

Thus, the confinement free energy becomes

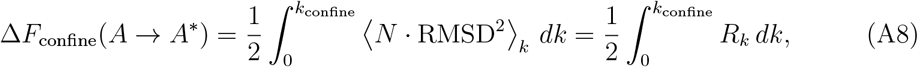

where 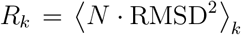. Therefore, by calculating the ensemble average of RMSD at different force constants *k*, we can estimate the confinement free energy through numerical integration.

For the integration, we adopt the numerical scheme proposed by Karplus and co-workers, which utilizes a trapezoidal rule on a double-logarithmic scale and assumes a power-law form *R*_*k*_ = *ak*^*b*^. The area under the curve between two successive points 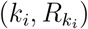 and 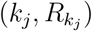, where *j* = *i* + 1, is given by

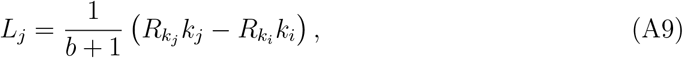

where *b* is calculated as

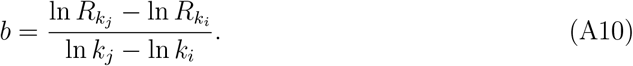

The total confinement free energy is then computed as the sum over all intervals:

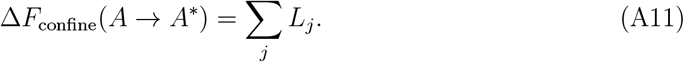

## Appendix B: Alchemical free energy for solvation/desolvation

To estimate the solvation or desolvation free energy of a protein, we employed an alchemical free energy perturbation approach. This method involves two thermodynamic states: the protein solvated in water (*A*^∗^) and the protein in vacuum 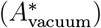. The transformation from the solvated to vacuum state (or vice versa) is performed by gradually decoupling water from the rest of the system using scaling parameters (*λ*). Non-bonded interactions (electrostatic and van der Waals) involving water molecules are parametrized by *λ* to remove their interactions with both themselves and other parts of the system. The total potential energy function is given by:

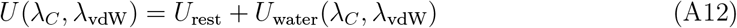

Here, *U*_rest_ is the potential energy of all components of the system excluding water while *U*_water_ accounts for interactions that involve water molecules. The alchemical transformation from vacuum to solvated state proceeds in two stages:

**Step 1 (Van der Waals coupling)**: Van der Waals interactions are gradually introduced while keeping electrostatics off:

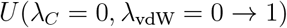

**Step 2 (Electrostatics coupling)**: Electrostatic interactions are turned on while van der Waals are fully present:

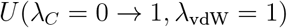

The total free energy change for the solvation is computed as sum over all intermediate *λ* windows:

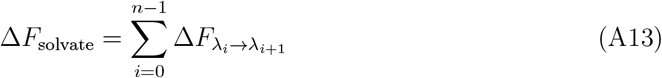

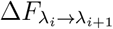 is calculated using the Bennett Acceptance Ratio (BAR) method which provides an optimal estimator based on bidirectional sampling:

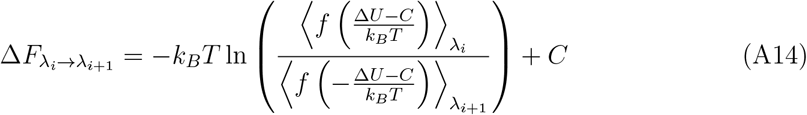

where 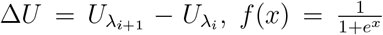 is the Fermi function, *C* is a constant solved self-consistently as

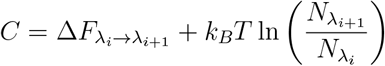

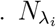 and 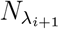 are the number of independent configurations sampled from the two states.

## Appendix C: Free energy from Quasi-Harmonic Approximation

The absolute free energy of the confined states in vacuum (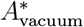 and 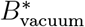) can be estimated using the quasi-harmonic approximation (QHA). QHA assumes that the system fluctuates within a harmonic-like, quadratic potential well, which should result in Gaussian distribution of atomic fluctuations. Under this approximation, the variance–covariance matrix of atomic fluctuations is computed as:

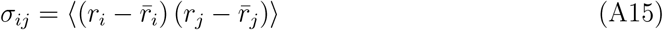

From this, a 3*N* × 3*N* mass-weighted Hessian matrix is constructed, where the effective force constant-like elements are given as:

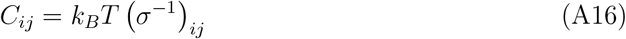

The generalized eigenvalue equation can then be written as:

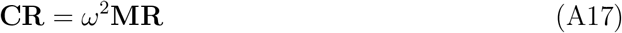

This can be reformulated using mass-weighted coordinates as:

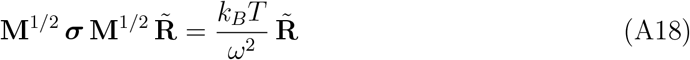

Here, **M** is the mass matrix, **R** is the displacement vector, 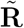is the mass-weighted displacement vector, and *ω* is the quasi harmonic frequency. The above form represents an eigenvalue problem, where 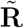 is the eigenvector and the corresponding eigenvalue is 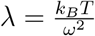.

In our study, all atoms in the protein are strongly restrained eliminating translational and rotational modes. Thus, only the 3*N* − 6 vibrational modes are used to estimate the vibrational entropy in the confined vacuum state:

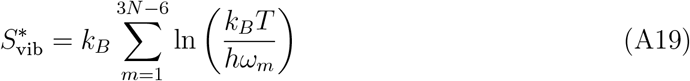

The absolute free energy of the confined state in vacuum, 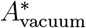(or 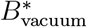), is given by:

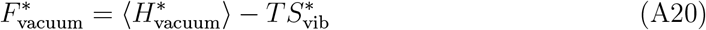

where 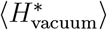 is the ensemble-averaged potential energy in vacuum.

## Acknowledgement

SP gratefully acknowledges GITAM for the Research Seed Grant (RSG/2024/0362) and access to computational resources through the High-Performance Computing facility, G-Cluster. SP also thanks Dr. Irawati Roy for valuable discussions on forcefield performance. AS and SP also thank Ashok Sekhar for his critical comments on this work. AS acknowledges the financial support from the Indian Institute of Science (IISc) and the high-performance computing facility "Beagle" that was set up from grants by the erstwhile IISc-DBT partnership programme. AS thanks the DST for the National Supercomputing Mission grants (DST/NSM/R&D-HPC-Applications/2021/03.10, DST/NSM/R&D-HPC-Applications/Extension Grant/2023/27). AS also acknowledges the FIST program sponsored by the Department of Science and Technology, India that supports the MBU infrastructure. AS would also like to thank the Teams Science Grant from the DBT-Wellcome Trust India Alliance (Grant number: IA/TSG/21/1/600245). AS also thanks the DBT National Network Project (NNP) grant (BT/PR40323/BTIS/137/78/2023) and the Matrics grants (MTR/2023/001040) from the Science and Engineering Board (SERB), India.

## Author contributions

AS and SP conceptualized the project. SP and AS designed the research. SP performed the research and analyzed the data. AS supervised the study. SP prepared the first draft of the paper. AS polished the draft.

## Data availability statement

Our code, models, and curated datasets are publicly available at the Zenodo repository [Repository-Thermodynamics-FoldSwitchingProteins](https://zenodo.org/uploads/17230657).

## Supporting Information

### Selection of forcefield

For metamorphic proteins, selecting an appropriate force field is crucial. We therefore tested three options: Amber ff99SB-ILDN^1^ with TIP3P water, Amber-disp,^2^ and CHARMM36 with TIP3P*.^3^ Using a seeding strategy in REMD simulations (Figure S1) as described by Parui et al.,^4^ we assessed the relative stability of the alpha and alpha/beta folds across different temperatures by measuring the population of both conformations. See Figure for more details. However, among the three, only CHARMM36 has shown the correct stability order within the physiological temperature range. We therefore selected CHARMM36 as our working force field and proceeded with the free energy simulations (CDCSR).

Amber ff99SB-ILDN: We first evaluated the Amber ff99SB-ILDN force field in combination with TIP3P water, using REMD in the temperature range of 278–370 K. At the lowest replica (278 K), this force field produced a higher population of *α/β* compared to 3*α* (Figure S2), which contradicts experimental findings where 3*α* should dominate at low temperature. Therefore, this force field was not considered further.

Amber-disp: The amber-disp force field was developed with the goal of providing a universal forcefield applicable to both folded and disordered proteins. ^2^ To evaluate its performance, we first examined the low-temperature population trends using REMD simulations in the range of 278–335 K. At the lowest replica (278 K), amber-disp stabilizes the 3*α* state (data not shown), which is consistent with the experimentally observed preference at low temperature. To assess the high-temperature behavior, we performed an additional REMD simulation starting at 340 K. Surprisingly, even at such high temperature, 3*α* remained more populated than the *α/β* state (Figure S3), contrary to experimental expectations where *α/β* should dominate. Due to this discrepancy, the amber-disp force field was excluded from further consideration.

CHARMM36: We next evaluated CHARMM36 with the TIP3P* water model (see Figure S4). To assess the low-temperature behavior, we analyzed the REMD with lowest replica at 278 K. At this temperature, the 3*α* conformation shows the highest population, consistent with experimental observations at low temperatures. To examine the high-temperature regime, we performed another REMD simulation starting at 320 K. At this temperature, the *α/β* population exceeds that of 3*α*, again in agreement with experimental results at higher temperatures. Since CHARMM36 reproduces the correct stability trends across physiological temperature ranges more reliably than the other two force fields tested, we selected CHARMM36 as our active force field for subsequent CDCSR simulations and further analyses.

**Figure S1:**
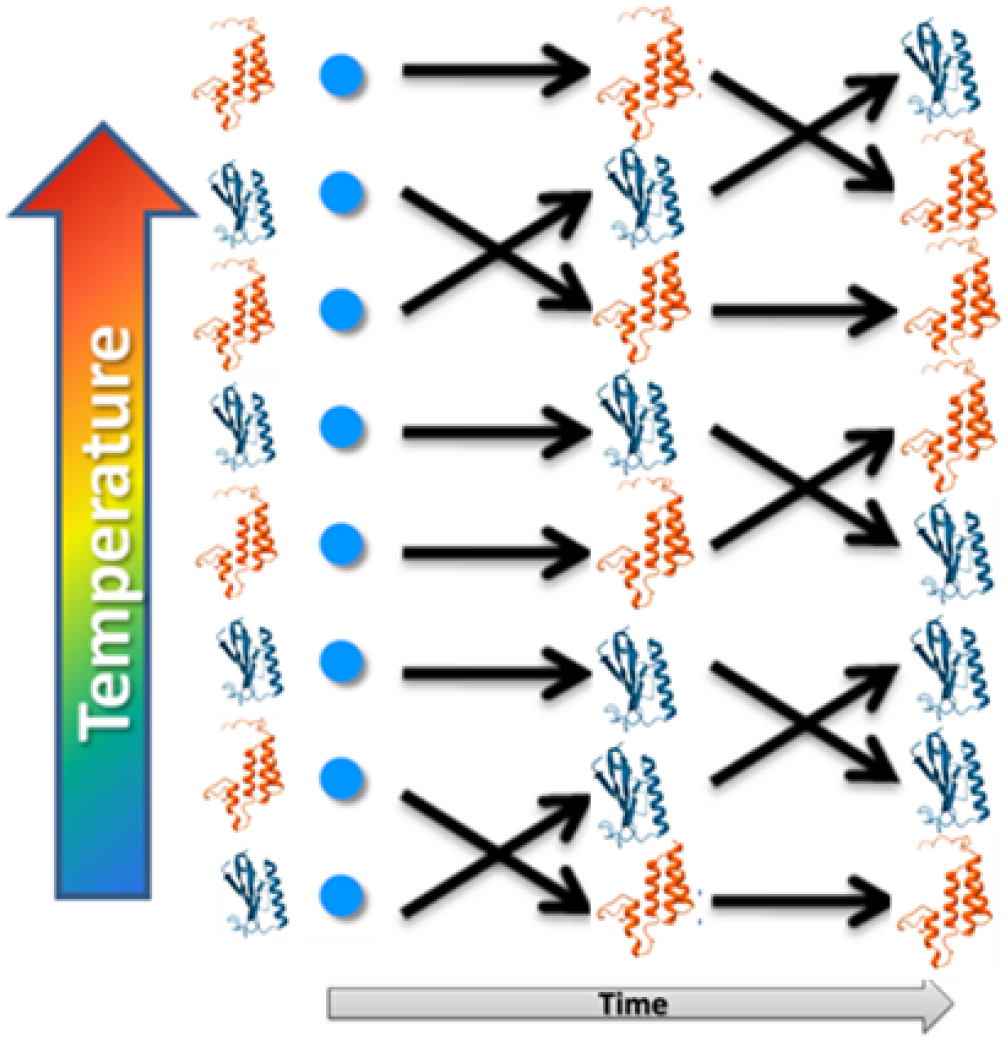
Schematic of a quick REMD-based strategy to probe the relative stability of 3*α* and *α/β* conformers. Equilibrated 3*α* and *α/β* structures are alternately seeded across the replica ladder (i.e., *α/β*, 3*α, α/β*, 3*α*, …). Within a few hours, the conformer populations at the lowest-temperature replica can be estimated. These populations provide a qualitative measure of relative stability. Appropriate force field was selected if the observed population trends aligned with the experimental trends that 3*α* dominates at low temperature, while *α/β* is favored at higher temperature. Although this approach does not produce quantitative free-energy differences, it provides a fast and efficient means to assess force-field performance.

**Figure S2:**
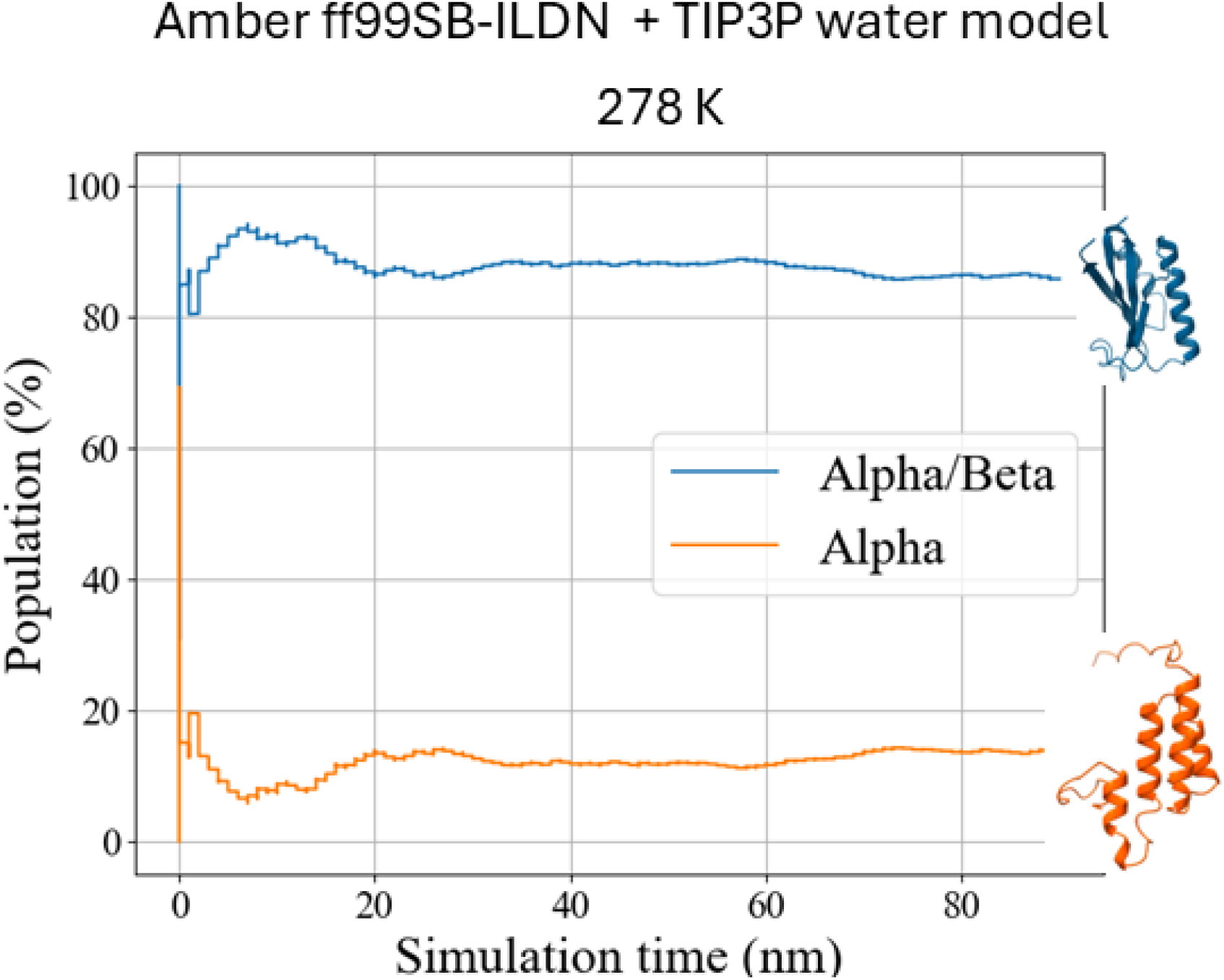
Populations of 3*α* and *α/β* at the lowest replica (278 K) for Amber ff99SB-ILDN with TIP3P water. Although 3*α* is experimentally more stable at this temperature, the force field instead stabilizes *α/β*.

**Figure S3:**
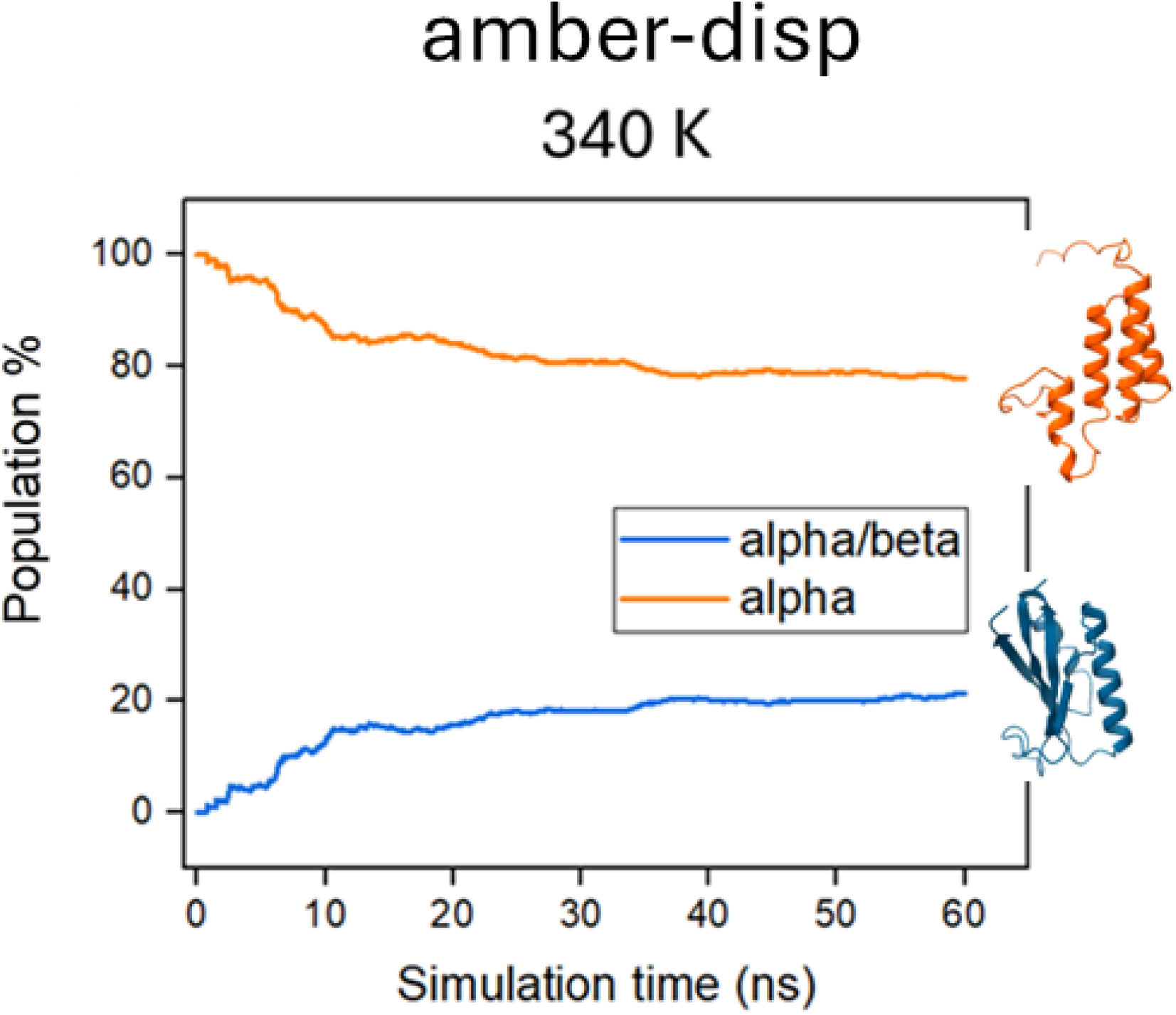
Populations of 3*α* and *α/β* at 340 K for amber-disp. Although experiments indicate that *α/β* is more stable at this temperature, amber-disp instead favors 3*α*.

**Figure S4:**
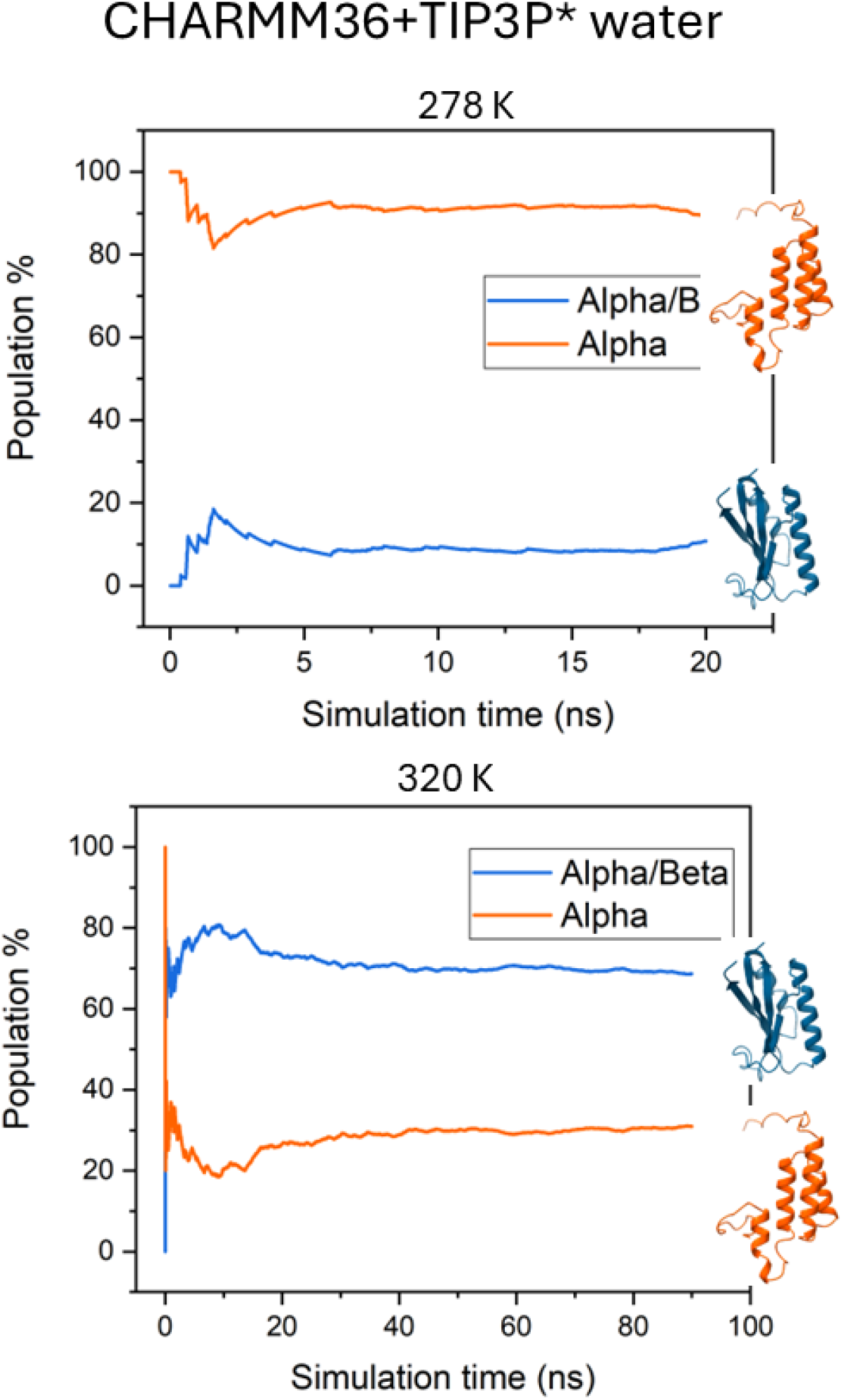
Populations of 3*α* and *α/β* at 278 K and 320 K obtained with CHARMM36. The force field captures the experimentally observed stability trends at both low and high temperatures.

**Table S1:**
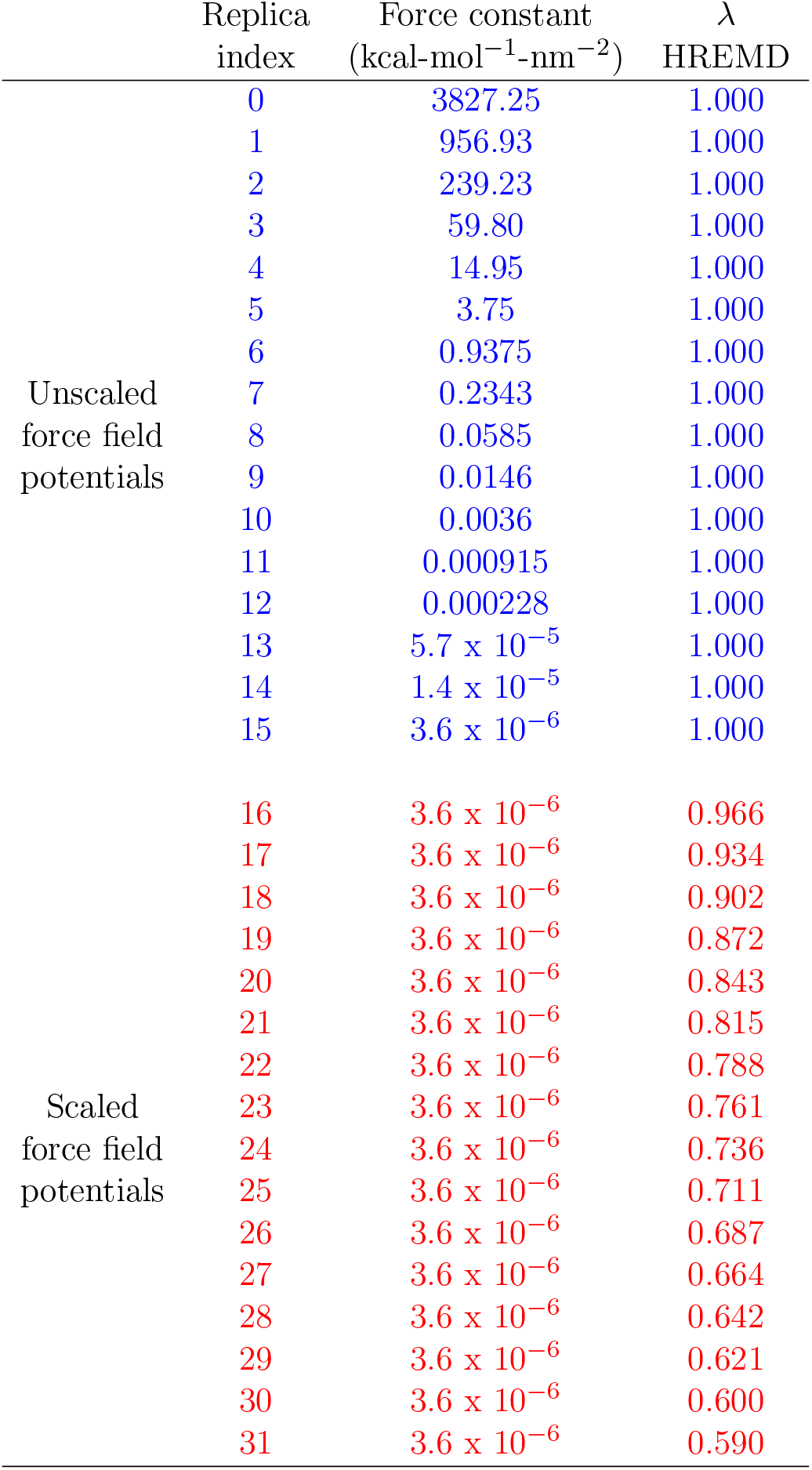
Confinement and Release Simulations. Replica 0 represents the fully confined state, while Replica 15 corresponds to the free state. Enhanced sampling is achieved by perturbing the force field beyond Replica 15 (red). The confinement and release free energy is calculated by analyzing transitions from Replica 0 to Replica 15 (blue).

**Table S2:** Solvation and Desolvation Simulations. Scaling (λ) values used in different replicas are shown. Replica 0 corresponds to the fully scaled state (λ = 0.0), representing the vacuum state. Replica 15 is the unscaled state (λ = 1.0), corresponding to the fully solvated system. λ for van der Waals potentials linearly increases from 0.00 to 1.00 across replicas 1 to 7 and remain constant at 1.00 for replicas 8 to 15. Coulombic potentials remain fully scaled (0.00) up to replica 7, after which coul-*λ* increases linearly from 0.00 to 1.00 across replicas 8 to 15.

| Replica index | vdw-λ | coul-λ |
| --- | --- | --- |
| 0 | 0.00 | 0.00 |
| 1 | 0.15 | 0.00 |
| 2 | 0.30 | 0.00 |
| 3 | 0.45 | 0.00 |
| 4 | 0.60 | 0.00 |
| 5 | 0.75 | 0.00 |
| 6 | 0.90 | 0.00 |
| 7 | 1.0 | 0.00 |
| 8 | 1.00 | 0.15 |
| 9 | 1.00 | 0.30 |
| 10 | 1.00 | 0.45 |
| 11 | 1.00 | 0.60 |
| 12 | 1.00 | 0.75 |
| 13 | 1.00 | 0.90 |
| 14 | 1.00 | 1.00 |
| 15 | 1.00 | 1.00 |

**Figure S5:**
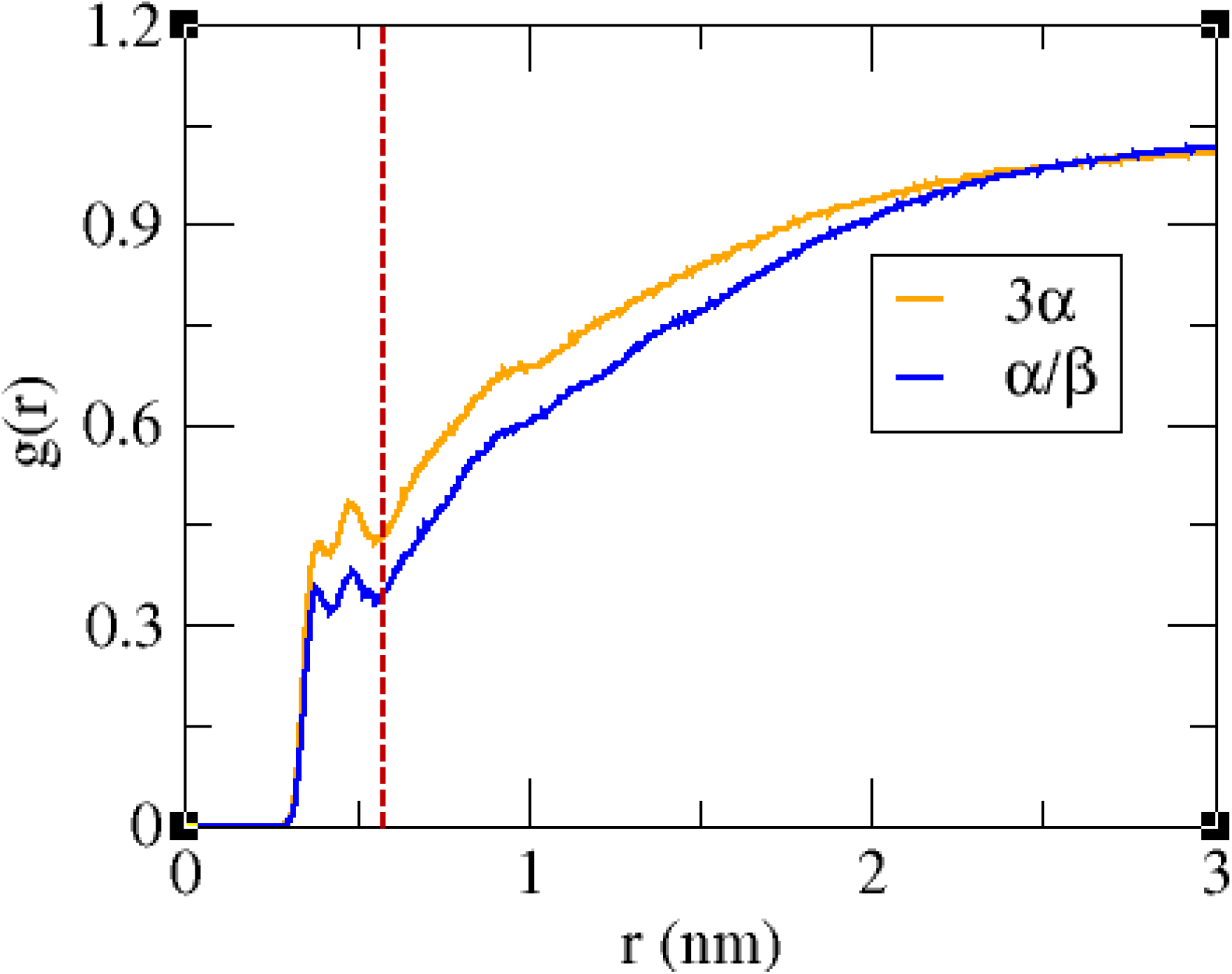
Radial distribution function of water (considering oxygen atoms) from the C-alpha atoms of protein. First solvation shell (FSS) of protein is designated by red dootted line (0.55 nm). Density of water in FSS of 3*α* is higher than that of *α/β*.

**Figure S6:**
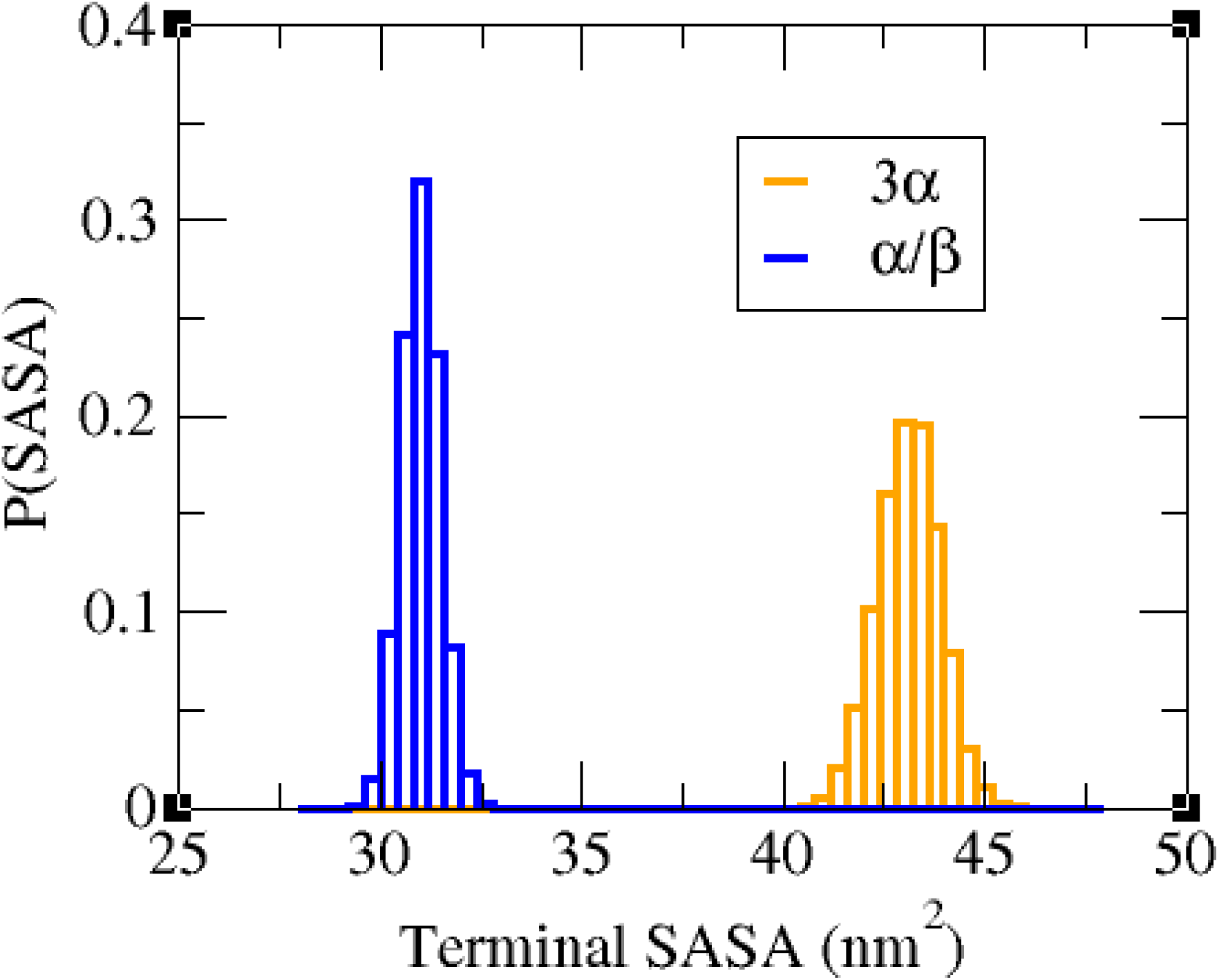
Distribution of solvent accessible surface area (SASA) of terminal regions (1-15, 83-94) of protein in both 3*α* and *α/β* form. Terminals of 3*α* form are more solvent exposed.

**Figure S7:**
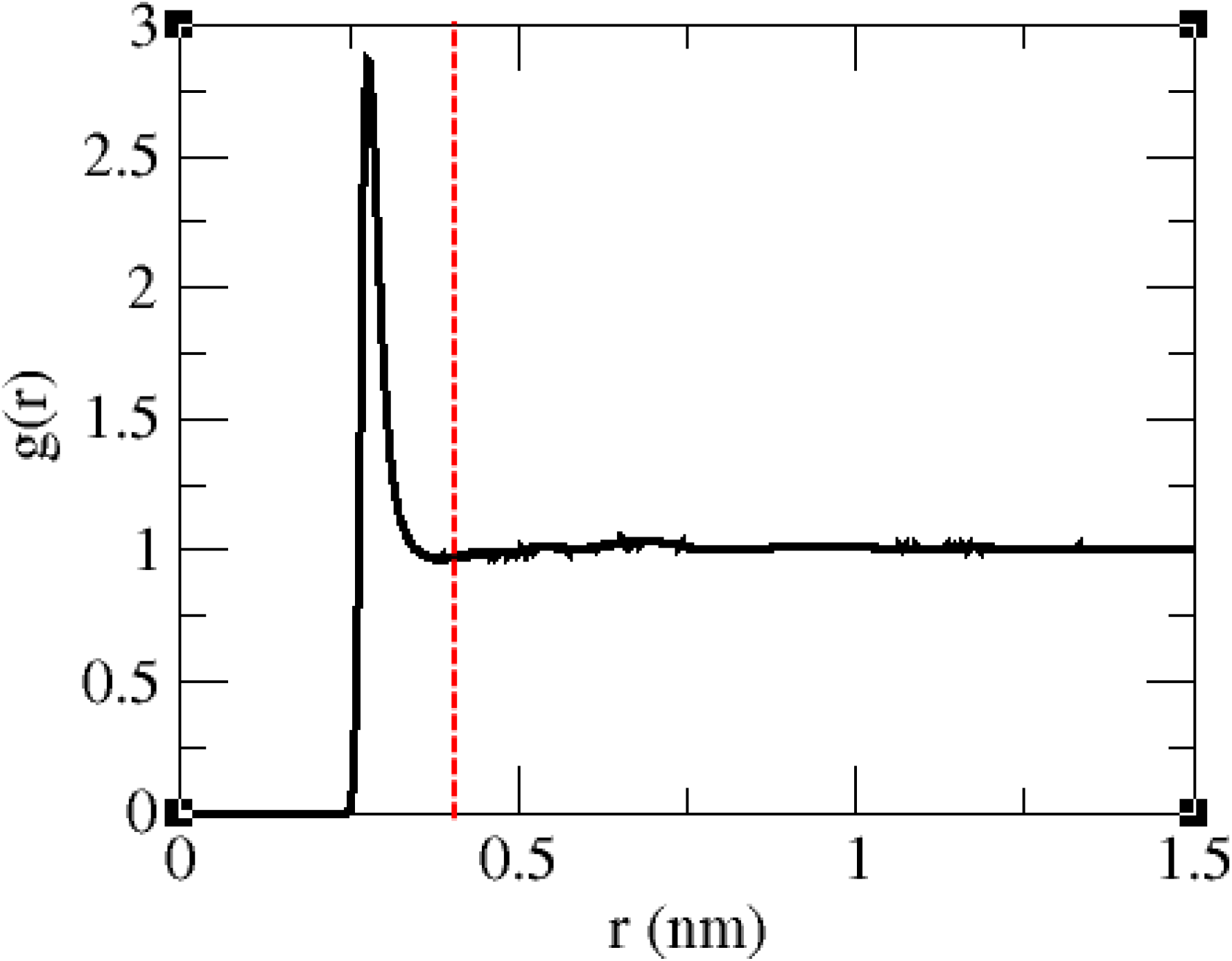
Water-water radial distribution function. Red dotted line highlights the first layer (0.4 nm) of water from the central water.

**Figure S8:**
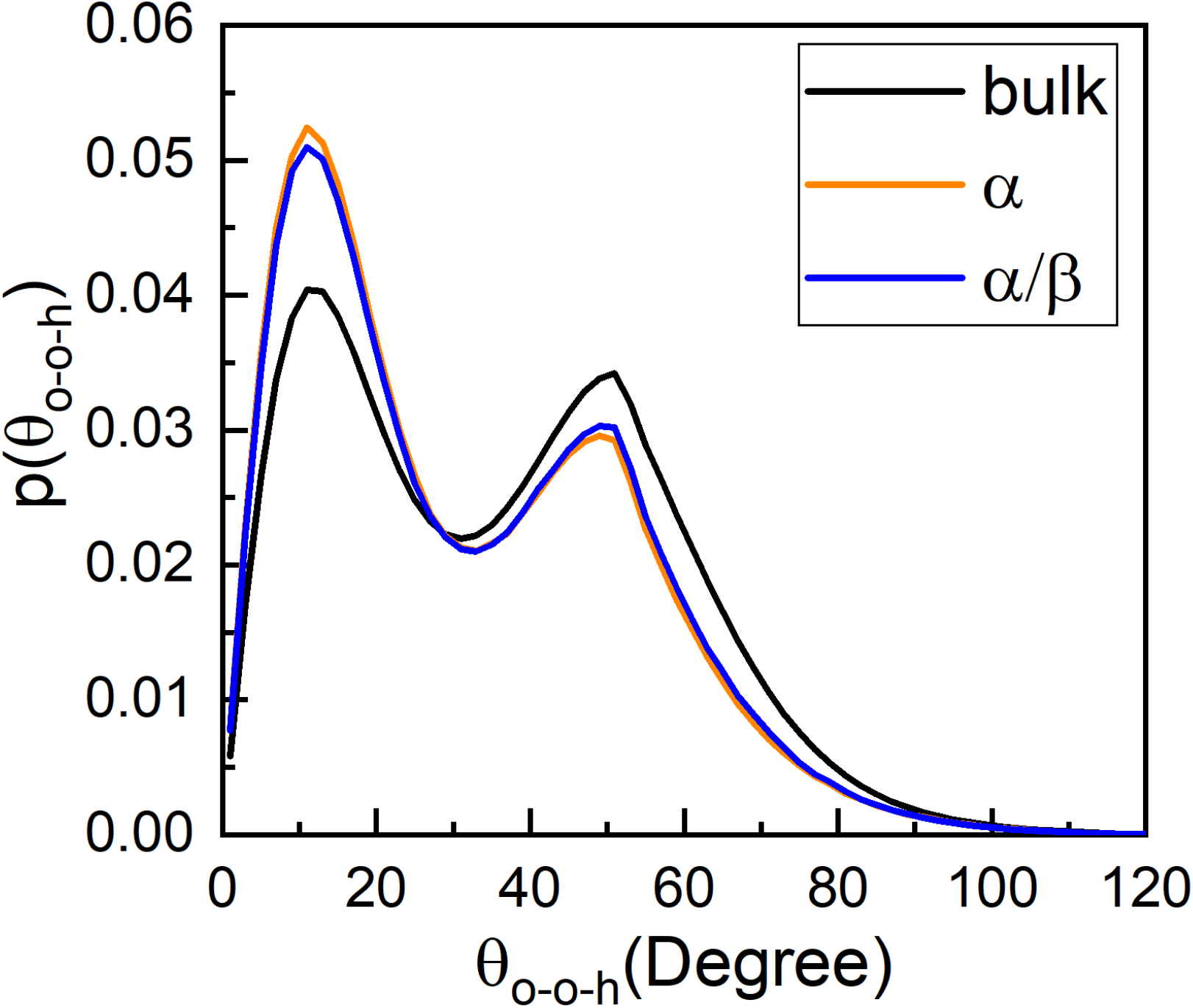
O-O-H angle distribution of water pairs (if *r*_*O*−*O*_ *<* 0.4 nm) in bulk, FSS of 3*α* and *α/β*. Distribution is composed of two peaks: peak at ∼ 15^*o*^ corresponds to stronger linear hydrogen bond and peak at ∼ 50^*o*^ is for weaker bent bond. Compared to bulk, water molecules around 3*α* and *α/β* are more strongly hydrogen bonded. Between the two folds, 3*α* has slightly higher population of stronger hydrogen bonds.

**Figure S9:**
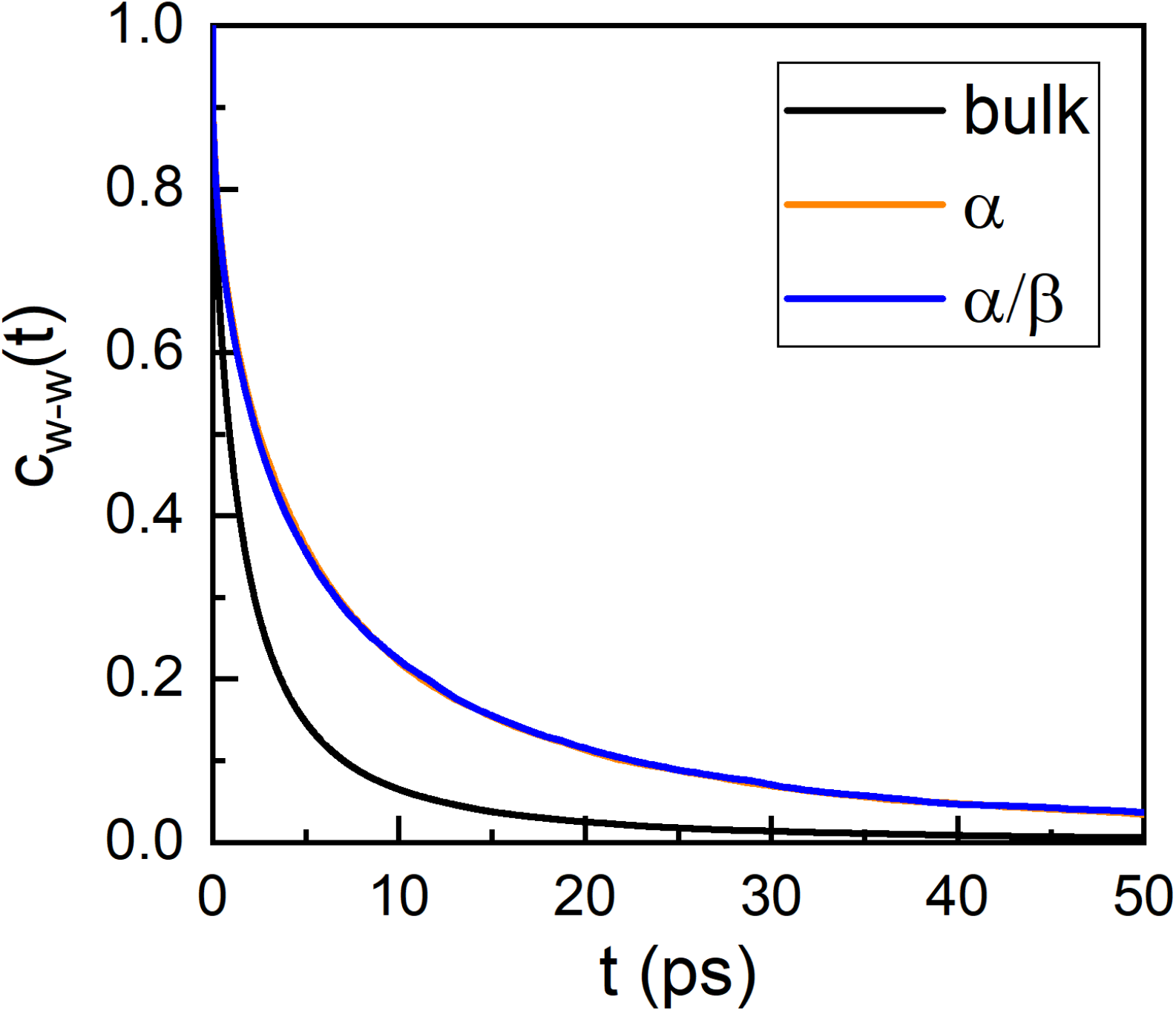
Autocorrelation function of water hydrogen bonds in bulk, FSS of 3*α* and *α/β*. Hydrogen bond lifetime of water around 3*α* and *α/β* is higher than that of bulk of water. However, decay rates in both 3*α* and *α/β* are almost similar.

**Figure S10:**
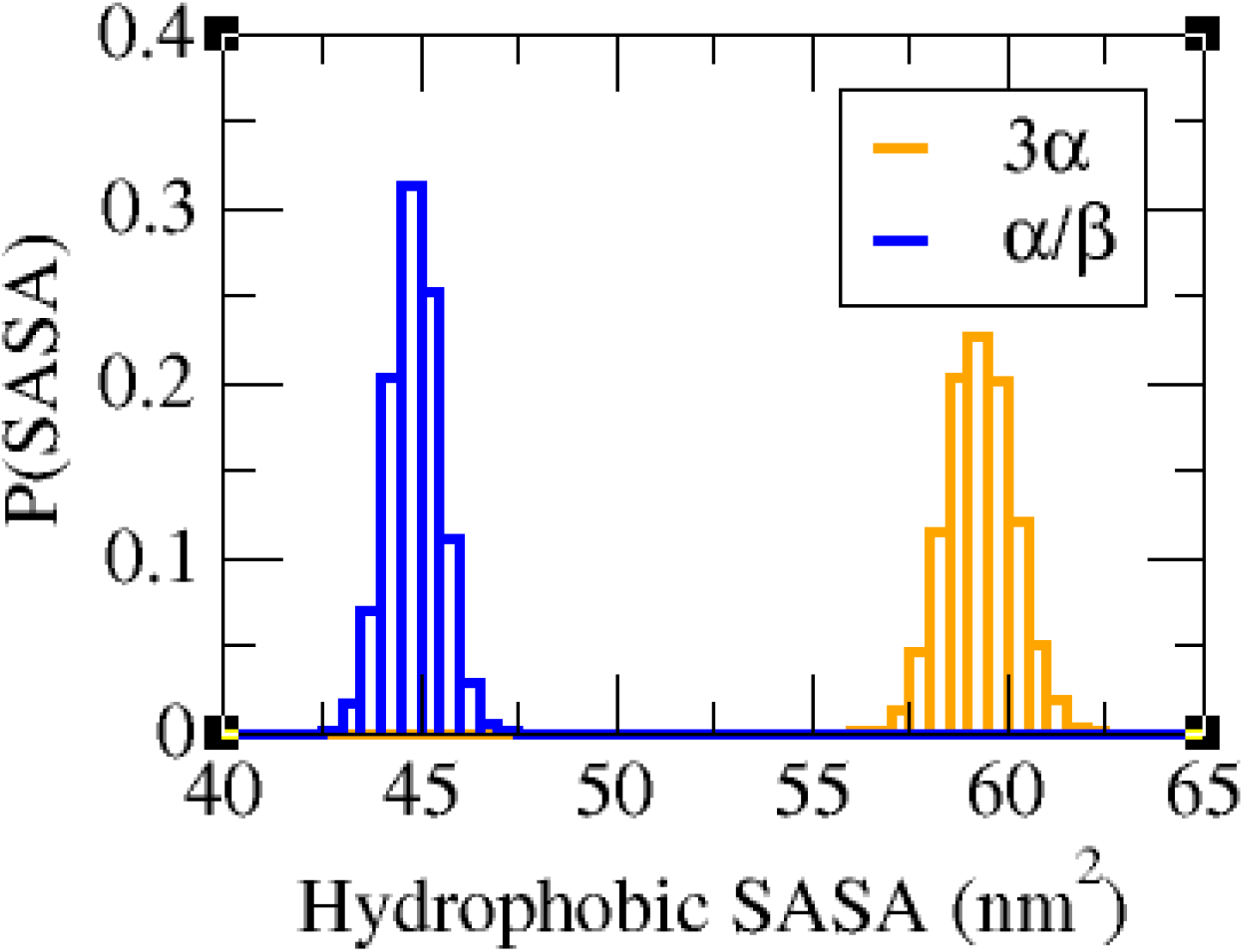
Solvent-accessible surface area (SASA) of hydrophobic residues in 3*α* and *α/β* folds. The *α/β* fold is more compact, whereas hydrophobic residues in 3*α* are more exposed to the solvent.

